# Animal group size variation in a minimal attraction-repulsion agent-based model

**DOI:** 10.1101/2024.03.20.585938

**Authors:** Jasper A.J. Eikelboom, Arjen Doelman, Frank van Langevelde, Henrik J. de Knegt

## Abstract

Grouping behaviour of prey animals is thought to be mainly driven by fear of predation and resource scarcity. Fear of predation often leads to small inter-individual distances, while resource scarcity leads to the opposite. Consequently, it is believed that the number of individuals in a group (group size) is an emergent property of the trade-off between acquiring scarce resources and preventing predation. We analysed whether group size can be reliably used as a proxy for this trade-off, using a deterministic attraction-repulsion agent-based model in a homogeneous area. In our model, each individual experiences distance-dependent attraction and repulsion to all others in the area, where varying degrees of grouping behaviour emerge from the number and distance of intersections between the attraction and repulsion functions. We show that the coefficient of variation of group size generally lies between 50 and 150%, depending on both animal density and the trade-off between resource scarcity and predation. Given that the variations of group size are already this large in homogeneous and deterministic scenarios, we urge researchers to be cautious in using group size as a proxy for the resources/predation trade-off and consider inter-individual distance as a more direct and potentially more reliable alternative.

## 1 Background

In order to survive, the movement of animals is for a large part driven by two factors: resource availability (e.g., food) and fear of threats (e.g., predation) [1, 2]. Resource scarcity induces animals to prioritize foraging behaviour and high predation risk induces animals to prioritize vigilance behaviour [3]. Given that these two behaviours often cannot be performed efficiently simultaneously, a trade-off exists between food acquisition and predator avoidance regarding optimal fitness [4]. Emerging from individual movement, collective animal movement (e.g., group formation) is to a large extent also shaped by both resources and predation [5, 6, 7]. In general, when the chance of predation is high it benefits an individual to live in a group with many individuals, through both the dilution (i.e., less chance to be chosen by a predator during an attack [8]) and the “many eyes” effect (i.e., benefiting from the vigilance of group members [4]). And although there are animal species that benefit from a sizeable group for resource acquisition (e.g., in order to defend territories with resources [5]: contest competition), when resources are accessible to all competitors (i.e., scramble competition) it benefits an individual to live solitary in the absence of predators by having the monopoly on resources in its direct vicinity [9]. Considering prey species that are in scramble competition with their conspecifics, there is thus an apparent trade-off between resource availability and fear of predation regarding an ‘optimal’ group size [5, 6].

The proposed mechanisms by which individuals form groups of a certain size are derived from two main theories: 1) individuals selecting groups of a certain size [10], and 2) individuals forming groups through self-organizing processes of individual movement decisions [11, 6]. Both mechanisms are credible to occur in nature, possibly even occurring simultaneously for some species [5]. However, the first of these two mechanisms is cognitively the most complex by assuming that animals are able to count conspecifics within a group, which is unlikely for large groups and/or species with lower cognitive abilities such as shoaling fish, swarming insects and flocking birds [6]. Furthermore, with the first mechanism, optimal group sizes are generally exceeded because solitary individuals continue to join already large groups [11]. Self-organization is thus the most parsimonious of these two mechanisms through which to explain the formation of group sizes that may appear optimal given the trade-off between searching for scarce resources and reducing predation risk [6]. These self-organizing models of collective behaviour have been build around individual movement decisions, in which often the radius of interaction with conspecifics and/or the magnitude of attraction and repulsion between individuals is a function of variables such as resource availability and fear of predation [11, 6, 12].

In many studies the concept of optimal group size has been used to draw conclusions about resource availability [13, 9], predation risk [14, 15] or both [16, 11] by monitoring group sizes in the field. However, group sizes are often highly variable, resulting in broad frequency distributions when recording the sizes of groups [13, 11]. This variability is not an issue in laboratory settings or during experiments where resource availability and fear perception can be controlled [11], but it does become problematic under field conditions where resource availability and predation risk are largely unknown and which are often the actual variables of interest for which group size serves as a proxy [13, 14]. To substantiate this, a large variance in field-monitored group sizes can be caused by 1) heterogeneity in resource availability or predation risk across the study area, 2) complexity in animal behaviour (which is obviously driven by many more factors), as well as 3) a large inherent variability in the emergent properties of the group formation process even in the absence of environmental and behavioural heterogeneity. To draw reliable conclusions about resource availability and predation risk from monitored group sizes it is thus important to investigate the relative importance of each of these three aspects in their shaping of the variation in group sizes.

The inherent variability of group sizes caused by the group formation process itself can be studied with a minimal agent-based model with two types of forces: attraction (driven by predation risk) and repulsion (driven by resource scarcity). In a hypothetical situation with fear of predation and unlimited resources it is most beneficial for animals to all stack in the same location (attraction-only), and in the situation without fear of predation and with limited resources it is most beneficial for animals to distribute themselves perfectly overdispersed across the area (repulsion-only). Attraction-repulsion agent-based models have been used widely in biology, mathematics and physics research to study clustering, both to investigate equilibria [17, 18, 19, 20, 21] and collective movement patterns [22, 23, 24]. From these modelling efforts it has become apparent that groups with alternate stable sizes can form in a single simulation from random initial locations, deterministic movement and homogeneous areas [17]. How the variation in stable group sizes exactly relates to the underlying model parameters is of interest for this study.

Here we aim to investigate the inherent variability of group sizes that result from a deterministic and homogeneous self-organizing group formation process driven only by resources and fear. Using this approach we can measure the variance in group sizes that are caused solely by the group formation process, thereby avoiding the effects of environmental heterogeneity and other factors that influence animal behaviour. Our aim is thus not to provide a single realistic model for group size variation, but a simple model to gauge the effect of the inherent variability in the group formation process. To this end we have built an agent-based model in which each individual experiences distance-dependent forces to all other individuals: attraction (driven by predation risk) and repulsion (driven by resource scarcity), each force modelled with only one parameter. From the converged simulations of these models we computed the variances in group size and linked these to the different values of attraction, repulsion and animal density. To sample the entire probability distribution of stable group sizes, we ran multiple iterations of each unique model with random initial locations of the individuals.

## 2 Modelling

Full details about the modelling are in the Methods section.

We modelled various numbers of individuals in one, two and three dimensions in areas of different sizes with an agent-based model. Each individual experienced both an attraction and a repulsion force to each other individual, for which the magnitude varied deterministically based on inter-individual distances. The net resultant vector of all these forces resulted in the movement of the individual. For every simulation the locations of the individuals were initiated with complete spatial randomness and the simulations ran until the locations converged to a stable position. The single-parameter function of the attraction force versus distance was chosen to be hump-shaped with a long tail: being low at close distances, high at intermediate distances and low at far distances. The single-parameter repulsion force function was chosen to decrease exponentially with distance: being high at close distances and low at far distances. These distance-dependent forces simulate the tendency of group-living prey animals to group together with conspecifics in the vicinity and to maintain a certain inter-individual distance [19]. We used the attraction parameter *a* as a proxy for predation risk and the repulsion parameter *r* as a proxy for resource scarcity and did not make assumptions about the functional relationship between predation risk versus *a* and resource scarcity versus *r* (other than being monotonically positive).

We chose our functions and underlying *a* and *r* parameters in such a way that the combination of the attraction and repulsion forces in a net attraction force (through subtracting the one from the other) led to three distinct scenarios of relationships (Figure 1): 1) when *r* had the upper hand over *a*, with the net attraction being negative for all possible distances; 2) when *r* and *a* were closer together in their influence on the system, with the net attraction being positive for a certain distance range and negative both before and after this range; and 3) when the balance between *r* and *a* is shifted even further, the net attraction being positive for all distances beyond a certain range. Based on the results of previous studies done on clustering with attraction-repulsion agent-based models [17, 18, 19], we expected the first scenario to result in overdispersed systems without group formation, the second scenario to result in systems where multiple groups can form, and the third scenario to result in systems that have the capacity to form one group with all individuals.

**Figure 1.**
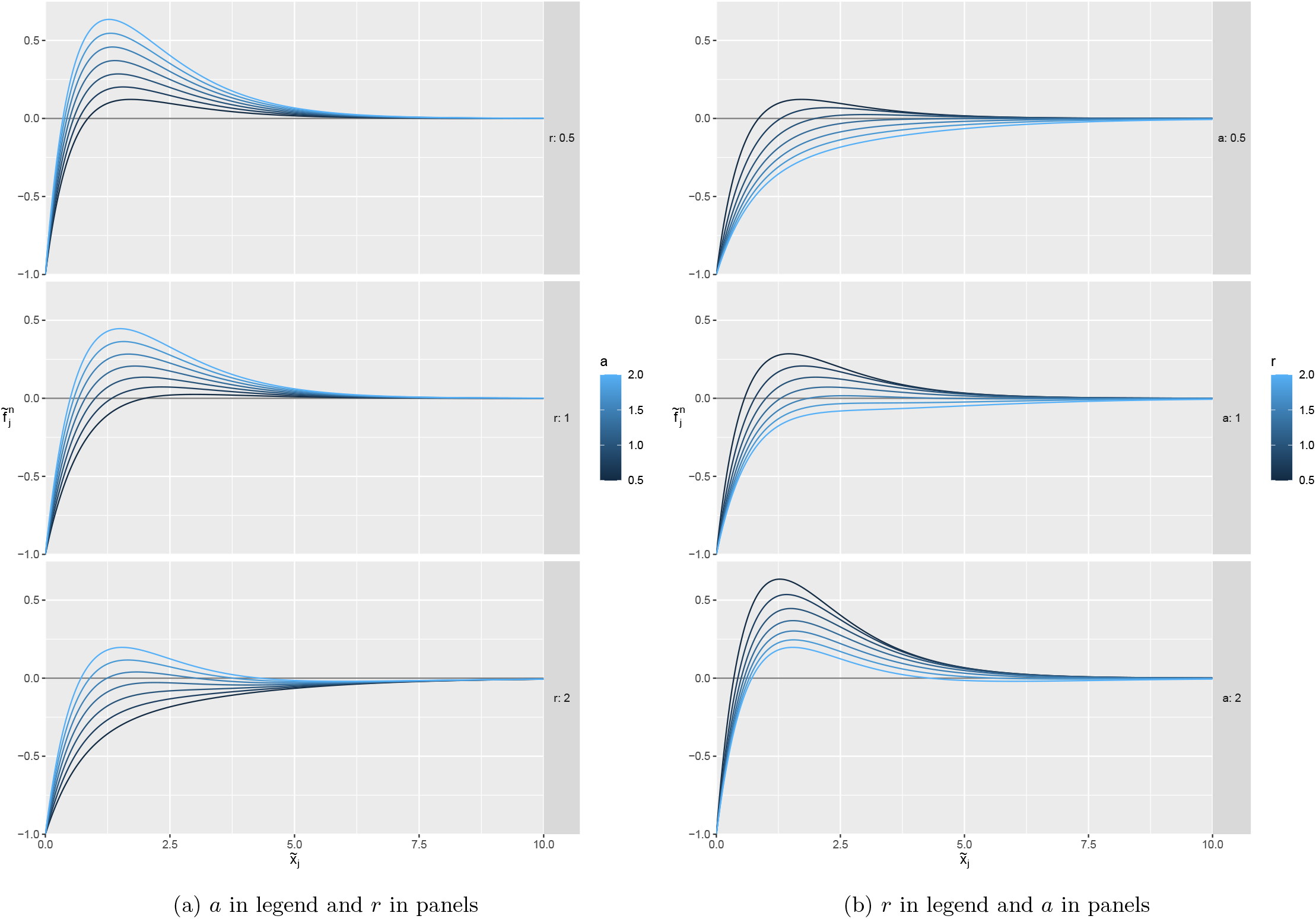
Net attraction 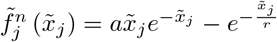 versus distance 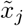, for values of attraction *a* and repulsion *r* between 0.5 and 2.

We simulated all parameter combinations of the model 100 times with different random initial locations of the individuals. In order to retrieve the group sizes of the converged states of the simulations, we first tried various clustering algorithms (e.g., *k* -means, *k* -medoids, hierarchical clustering and brute-force packages that applied many different clustering algorithms at the same time) to differentiate the individuals into different groups. Unfortunately this proved to be too error-sensitive for our data, especially for situations where the distances between clusters were not much larger than the distances between individuals within the same cluster. Therefore we computed the distance matrices of the individuals and scaled the inter-individual distances based on the expected inter-individuals distances under complete spatial randomness (see Clustering subsection). From these transformed distance matrices we then identified the values that were substantially lower than expected and used these to derive an accurate estimation of the distributions of group sizes.

## 3 Results

By visualizing the end locations of the simulations (e.g., see Figure 2), we noticed that group formation was as expected largely influenced by both the balance in *a* and *r* as well as the density of individuals. Unexpectedly though, many simulations formed separate multi-individual groups instead of the expected single “supergroup” when the net attraction relationship did not became negative again after a certain distance range. Furthermore, there were simulations which were completely overdispersed into solely 1-individual groups, even when the net attraction was positive for a certain distance range. This is likely because the size of the simulated area extends on purpose far beyond the effective distance range of the net attraction function to resemble realistic natural scenarios and because the density of individuals is for certain simulations relatively low.

**Figure 2.**
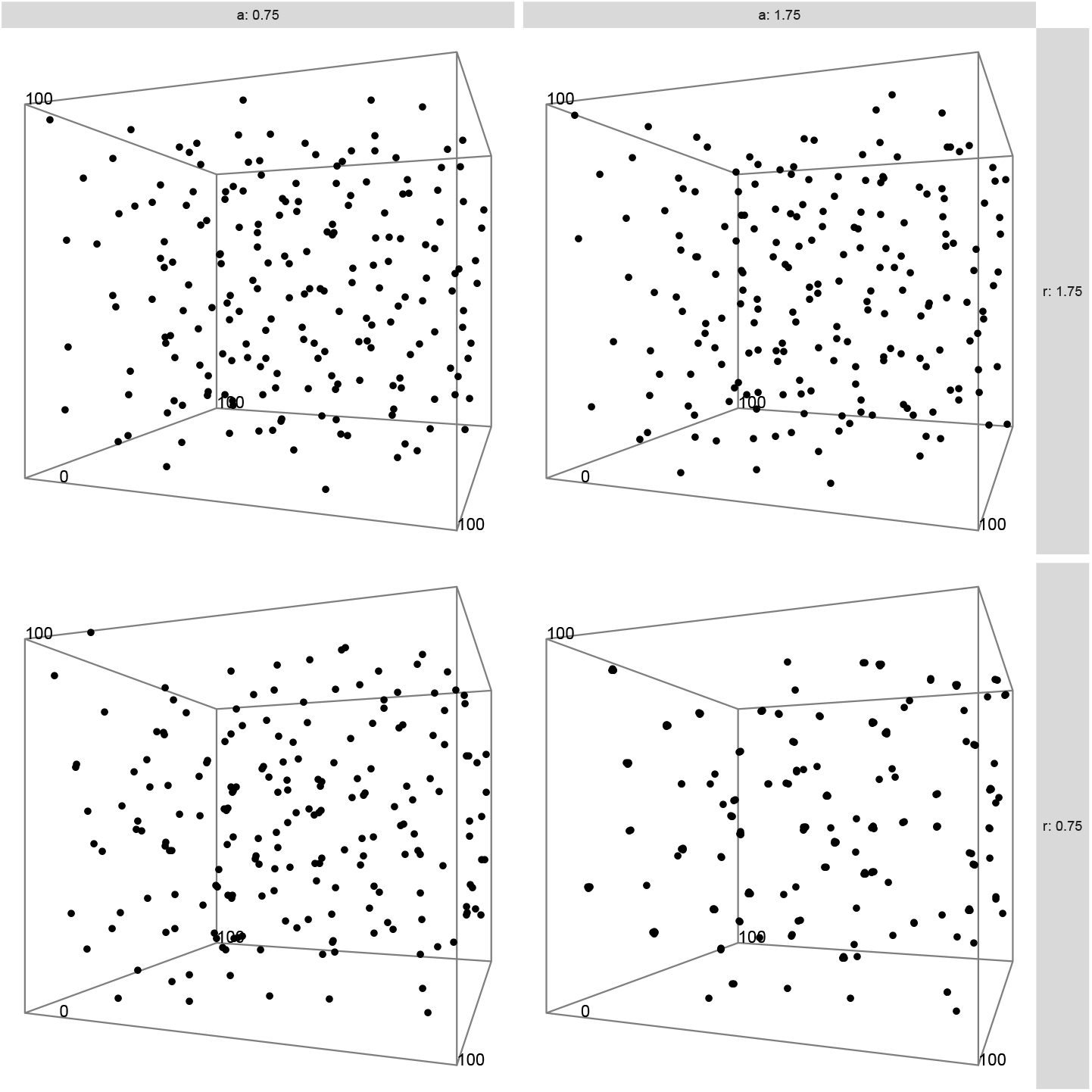
Converged states of 1 initialization of our model with 200 random initially distributed individuals inside a 3D toroidal landscape with a size *s* of 100, two different values of attraction *a* in the top panels and of repulsion *r* in the side panels. The two top visualizations display no clustering, the bottom left some clustering and the bottom right even more.

The transformed distance matrices allowed us to easily identify the individuals which were closer together than expected under complete spatial randomness (Figure 3), from which the distributions of *k*th neighbours within the same group and subsequently the probability distribution of an individual being in a group of a certain size could be derived.

**Figure 3.**
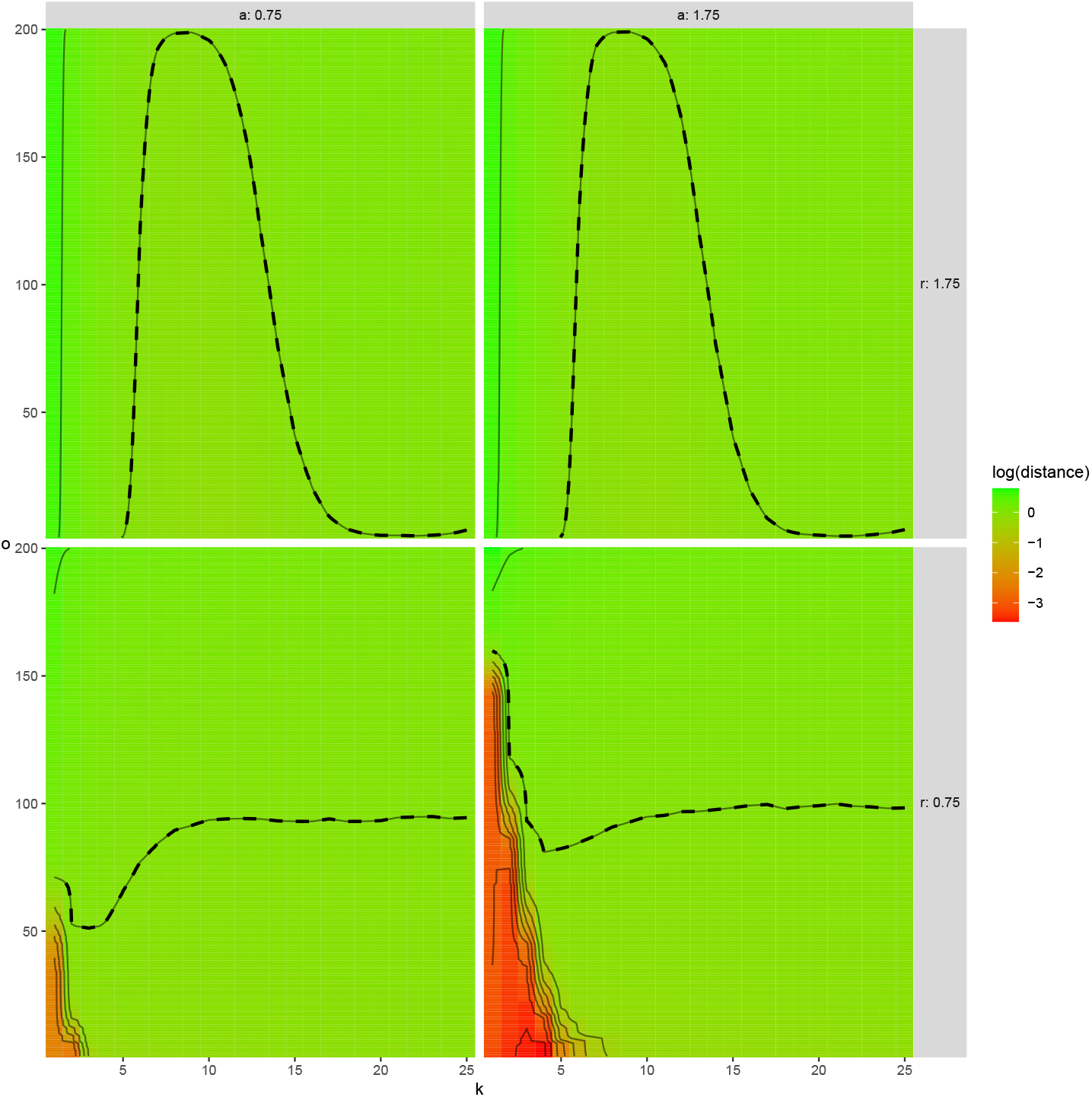
Average transformed distance matrices 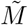 of the converged states of all 100 3D initializations as displayed in Figure 2, with the *k*th neighbour (until the 25th) on the x-axis, the ascending order *o* of distances on the y-axis, two different values of *a* in the top panels and of *r* in the side panels. The expected distance under complete spatial randomness is indicated with a dashed line. See Clustering subsection for details on the construction and use of these matrices.

After retrieving the distribution of group sizes for all parameter combinations of the model, we noticed that group size was as expected largely influenced by the difference between the distances for which the net attraction was zero (Figure 4, where the Lambert’s *W* value is a mathematical proxy for the difference between these distances).

**Figure 4.**
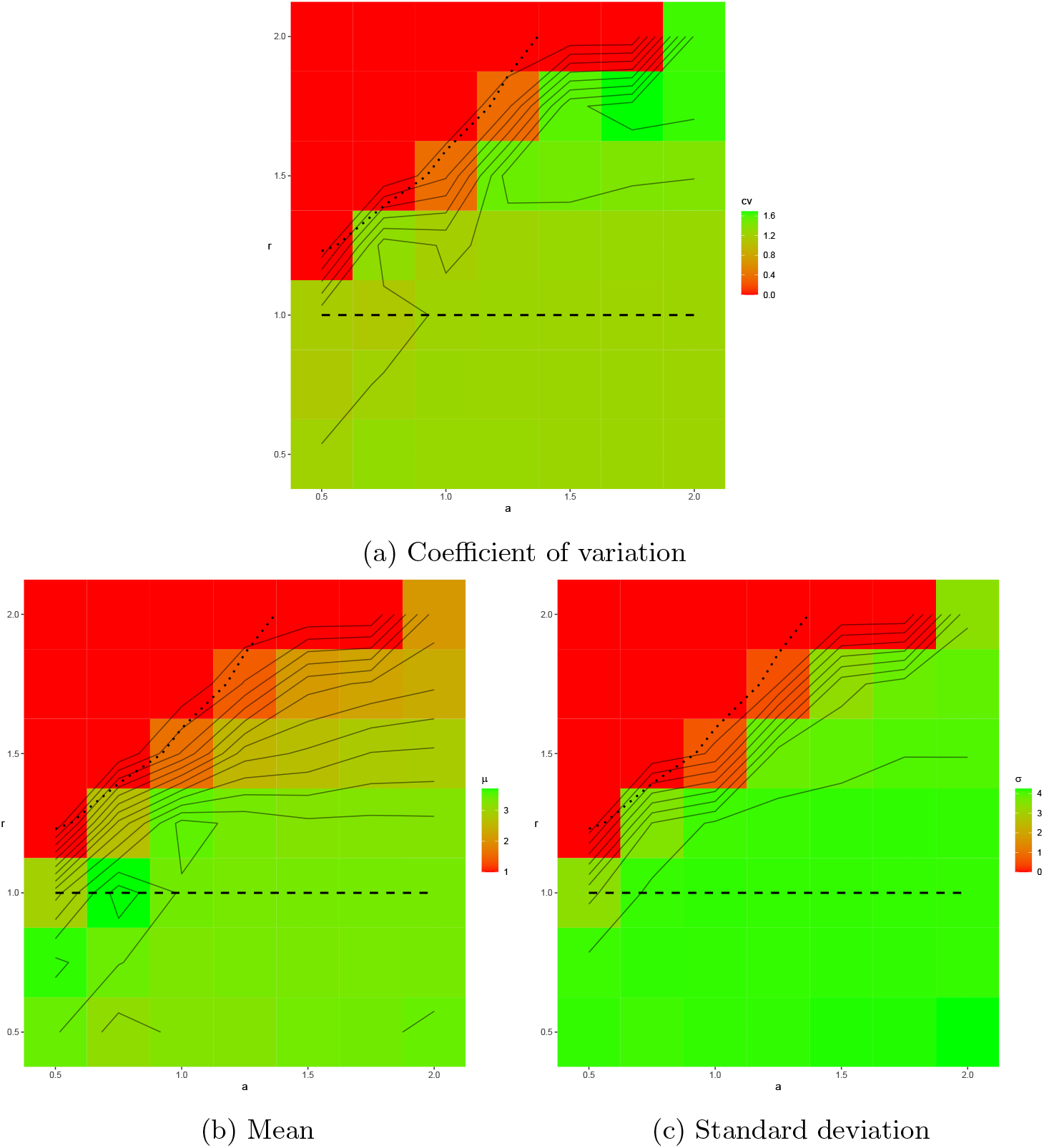
Group size versus attraction *a* (x-axis) and repulsion *r* (y-axis), for 1 dimension, 200 individuals and torus size 800. Coefficient of variation 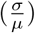 in Figure 4a, mean (*µ*) in Figure 4b and standard deviation (*σ*) in Figure 4c. The contour lines of 4b and 4c seem to follow the contour lines of the Lambert’s *W* value for the intersections between the attraction and repulsion functions (Figure 6), of which the dashed (indicating the threshold between 1 and 2 intersections) and the dotted line (threshold between 2 and 0 intersections) are visualized here as well. This indicates a relationship between the mean and standard deviation of group size with the distance between the intersections of the attraction and repulsion functions.

Given all unique combinations of the parameters of our simulations, the same relative patterns as in Figure 4 were produced, but with differences in absolute values of group size. We then aimed to combine the balance between *a* and *r* (as expressed in Figure 4) with all other simulated variables into two interpretable variables: 1) the mean proportion of the total number of individuals within the attraction range (defined as zero for a repulsion-only model and otherwise defined as the largest distance with a value one-tenth of the value of the peak of net attraction) of each individual during initialization, and 2) the ratio between the mean proportion of the total number of individuals within the repulsion range (defined as the distance where the net attraction first becomes positive) of each individual during initialization and the former variable. These two variables seemed to describe the mean group size relative to the total number of individuals well (Figure 5b), but for the variation in group size the total number of simulated individuals also had a large influence (Figure 5a and 5c). Overall, the mean group size increases linearly with the proportion of individuals within the attraction range and increases even further when the proportion of individuals within the repulsion range is low relative to the proportion of individuals within the attraction range (Figure 5b). The same applies to the standard deviation of group size, although this relationship does not seem to be increasing linearly, but seemed to increase quicker for lower values of the proportion within the attraction range (Figure 5c). Furthermore, with smaller numbers of individuals the standard deviation of group size is noticeably larger. Dividing the standard deviation by the mean value of relative group size results in the coefficient of variation (Figure 5c), which appears to be hump-shaped and leveling off for larger larger values of the proportion within the attraction range. The coefficient of variation of relative group size lies approximately between 50 and 150% for the various parameter combinations.

**Figure 5.**
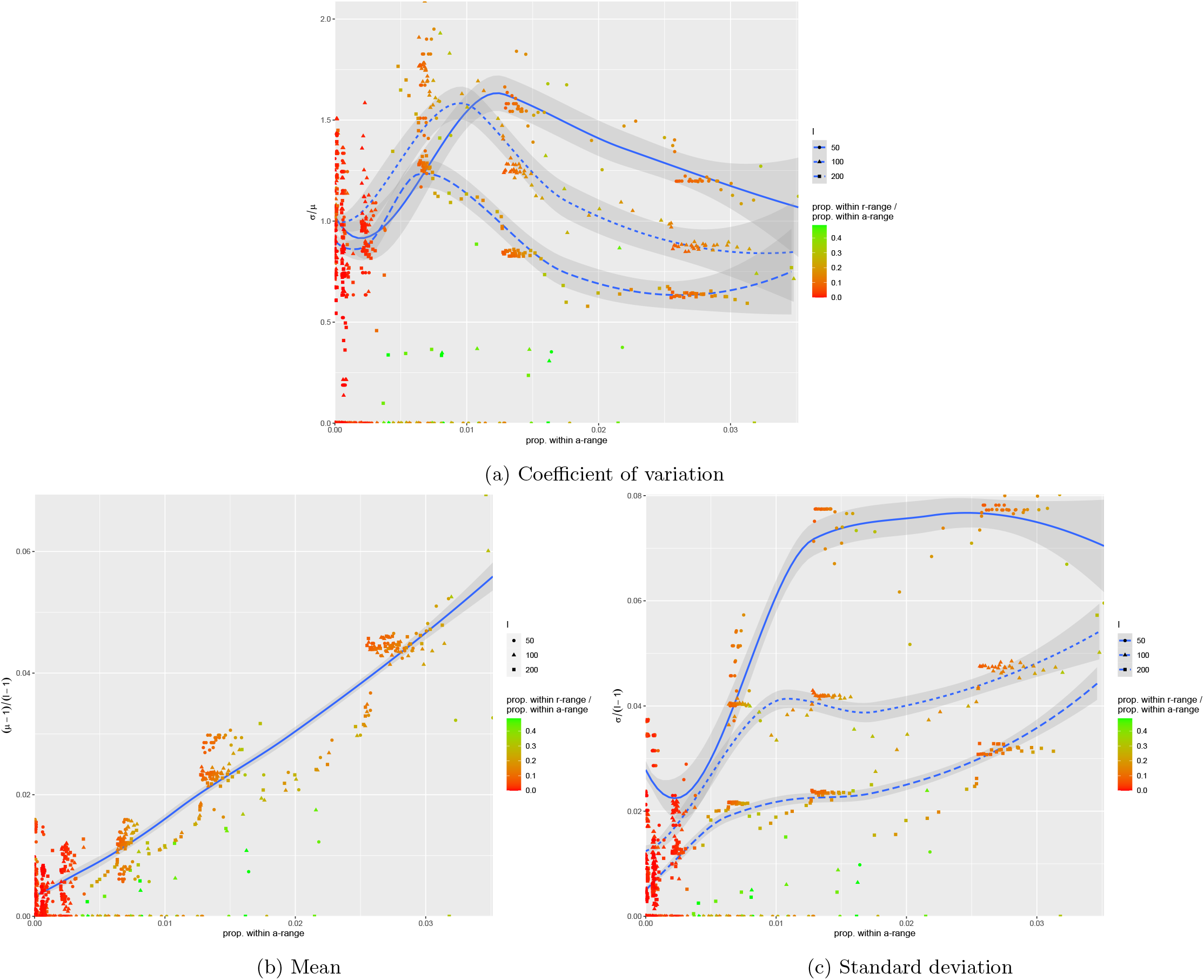
Group size proportional to number of individuals *l* (y-axis) versus average proportion of individuals under initial random distribution within every attraction range (x-axis), relative proportion within every repulsion range (color) and number of individuals *l* (shape). Coefficient of variation 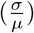 in Figure 5a, mean 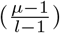 in Figure 5b and standard deviation 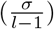 in Figure 5c. Trend lines computed independently of relative proportion within repulsion range and only when clustering occurred, using local polynomial regression fitting.

## 4 Discussion

In our study we have used an attraction-repulsion agent-based model to investigate the relationship between resource availability and predation risk versus animal group size. Using a single attraction parameter as a measure for predation risk and a single repulsion parameter as a measure for resource scarcity, we found that the mean relative group size increased with predation risk and resource availability, after having scaled the parameters for animal density. The standard deviation of relative group size behaved the same, but was noticeably larger for a smaller total number of animals and displayed a quicker increase for lower values of predation risk. As a result, the coefficient of variation of group size was highest for intermediate values of predation risk and depended on both the predation/resources trade-off and animal density. Overall, the coefficient of variation of group size generally lied between 50 and 150% in our simulations.

Our results show that mean animal group sizes increase with population density, predation risk and resource availability. Increased resource availability obviously allows for larger groups rather than causing it, as there is only a reason to cluster when there is a certain predation risk. Our results match well with empirical data, e.g., it has been shown that the mean group size increases with population density for fish [25], with predation risk for birds [15], resource availability for mammals [13], and all three factors for other taxa as well [5, 6]. Furthermore, an increase in predation risk translated to an increased range of attraction in our model, while a decrease in resource availability translated to an increased range of repulsion. This approach of interaction ranges has been used directly in other modelling studies, which yielded comparable results to our study regarding the probability distributions of group size. For example, a decrease in the local interaction radius of modelled fish resulted in a smaller mean group size as well as a smaller spread of the group size distribution [11].

The coefficient of variation of our modelled group sizes lies between 50 to 150% in homogeneous and deterministic scenarios, where the random initializations facilitated the observed variation. Given that there is such a large difference between the confidence limits of these group sizes, it is plausible that researchers who encounter such large differences in the field may actually interpret these as being caused by a difference in the predation/resources trade-off. There have for example been studies on ungulates [26], rodents [27] and monkeys [28] that have reported effects of a predation/resources trade-off on group sizes, while the variation in all group sizes was close to our reported variance here. This is potentially problematic, as the reported group sizes may thus have actually come from a single distribution (viz., with equal predation risk and resource availability).

Group compactness is a more direct proxy for grouping tendency than group size in situations where group formation is driven by local interaction rules, given that the distance between individuals follows directly from the interaction rules and group size is an emergent property of this self-organizing process [6]. For example, it has been shown that fish groups become more compact with increased predation risk [29]. Furthermore, group compactness seemed to be more important than group size in the preference of certain fish species [30], and group compactness also seemed more important than group size in the reduction of predation risk through a “confusion effect” [31]. However, group compactness has generally been used far less in the literature than group size to gauge the predation/resources trade-off, especially for terrestrial animals [9, 13, 14, 15, 26]. This is of course not surprising, given that it is easier to measure group size than group compactness, as compactness must preferably be monitored over a longer period of time and often requires equipment like cameras and tracking software while a single count suffices for group size. However, given the large inherent variability in group size given the same predation/resources trade-off, the usefulness and reliability of group compactness as a field proxy for this trade-off should be investigated for multiple species and study areas.

In this study we focused on purpose solely on the inherent variability of group sizes that result from a deterministic and homogeneous self-organizing group formation process, thereby leaving out other processes. Extra complexity in more realistic models can further amplify the variation in group sizes that we found (e.g., through local environmental heterogeneity or animal movement being dependent on more factors) or dampen it (e.g., forward persistence in movement that could lead to fission and fusion dynamics, which potentially leads to group sizes that become more ‘averaged out’ over time). It would therefore be recommendable for future research to also investigate the relative importance regarding the variability in group sizes of environmental heterogeneity and animal behaviour complexity in interaction with this simple group formation process.

## 5 Conclusion

We demonstrate self-organizing animal group formation with an attraction-repulsion agent-based model, for which the group sizes increase with predation risk, resource availability and population density. Even though this process is deterministic and homogeneous, the group sizes have a coefficient of variation between 50 and 150% depending on the aforementioned parameters. Such large variations in a single process are problematic when group sizes in the field are gauged on having resulted from differences in the predation/resources trade-off. We therefore urge researcher to investigate the usefulness and reliability of group compactness as a more direct proxy for the predation/resources trade-off.

## 6 Methods

### 6.1 Agent-based Model

Our agent-based model »is composed of *l* individuals of which each individual experiences attraction 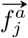 and repulsion 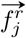 forces to all other conspecifics *j* based on their distance *x*_*j*_. For each point in time these forces sum up to one vector per individual 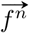, which determines the movement of that individual at that point in time.

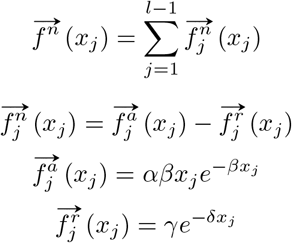

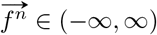, movement vector

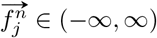, net attraction to conspecific *j* of *l* individuals

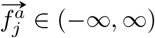, gross attraction to conspecific *j* of *l* individuals

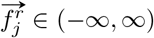, repulsion from conspecific *j* of *l* individuals

*x*_*j*_ ∈ [0, ∞), distance from conspecific *j* of *l* individuals

*α* ∈ (0, ∞), attraction height parameter

*β* ∈ (0, ∞), attraction rate parameter

*γ* ∈ (0, ∞), repulsion height parameter

*δ* ∈ (0, ∞), repulsion rate parameter

### 6.2 Nondimensionalization

Given that our model is not specifically designed for a certain animal species, the absolute values of *x*_*j*_ and 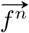 have no inherent meaning. Therefore, both dimensions of the net attraction function 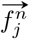 can be scaled to relative dimensions. This results in a simpler scaled net attraction function 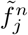 with two instead of four parameters.

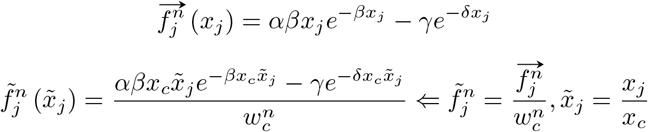

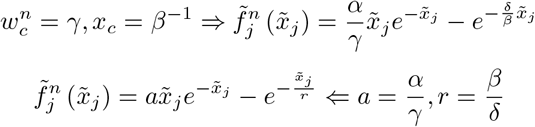

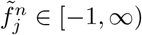, scaled net attraction to conspecific

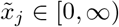, scaled distance from conspecific *j*

*a* ∈ (0, ∞), scaled attraction parameter

*r* ∈ (0, ∞), scaled repulsion parameter

### 6.3 Intersections

The intersections between the attraction and repulsion functions, i.e., when 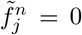, determine at which distances to a conspecific an individual remains stationary in the absence of other conspecifics. With multiple conspecifics, these intersections will determine the distances between conspecifics within the same group and the distances between groups. Note that the solution is a Lambert’s *W* function, which can have zero, one or two solutions (Figure 6).

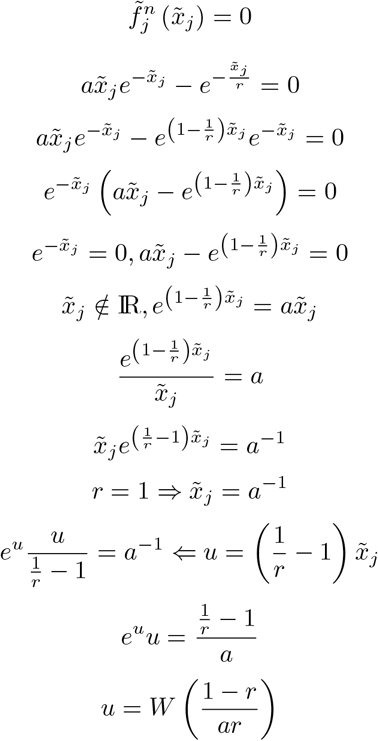

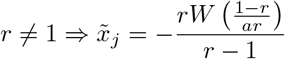

**Figure 6.**
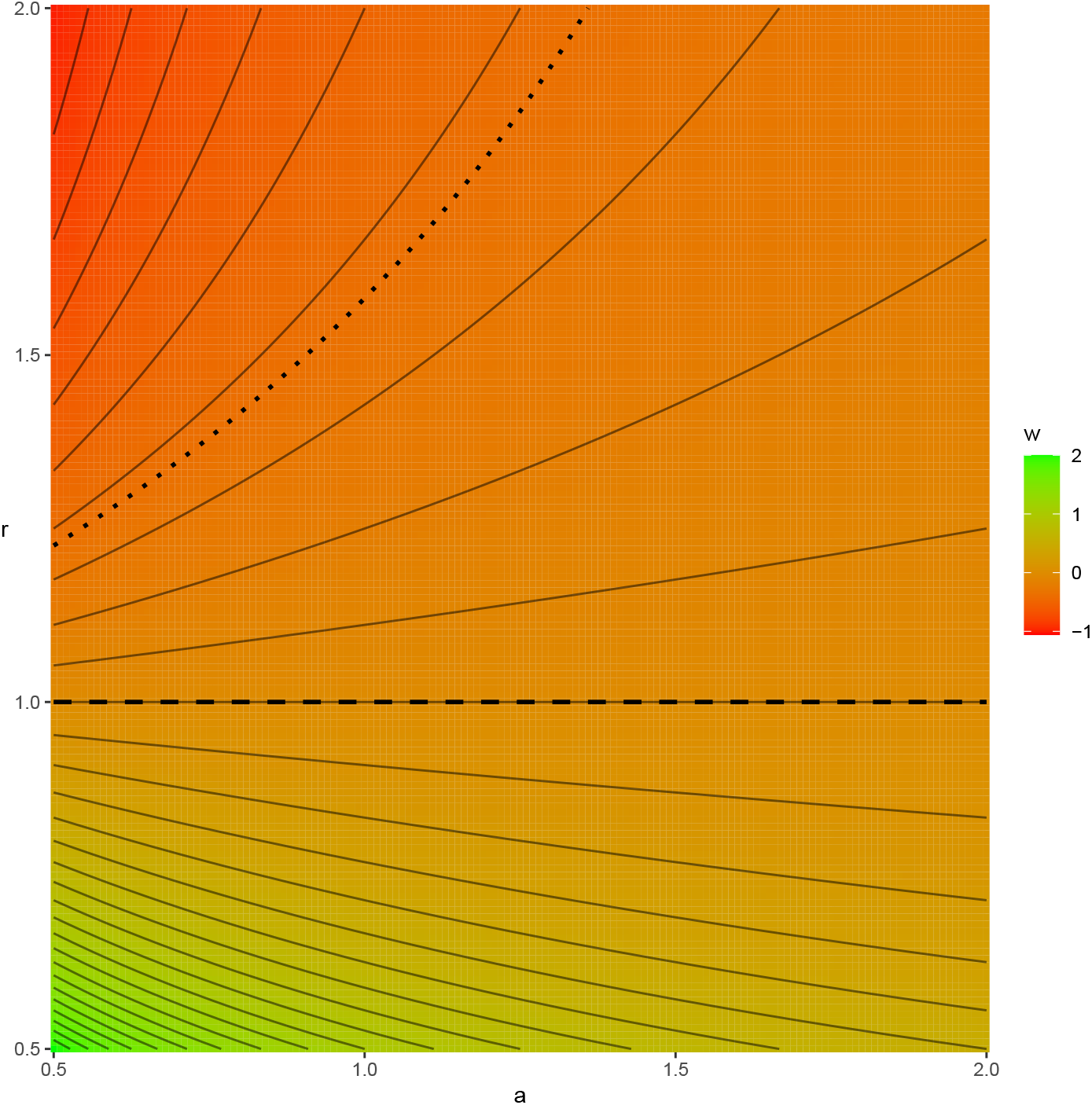
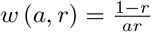, the input value of Lambert’s *W* function for intersections. *w* higher than 0 (dashed line) gives 1 intersection, *w* between 0 and −*e*^−1^ (dotted line) gives 2 intersections, and *w* lower than −*e*^−1^ gives 0 intersections.

### 6.4 Peaks

Following the same equation solving techniques as for the intersections, the peaks of 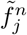 can also be derived.

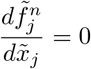

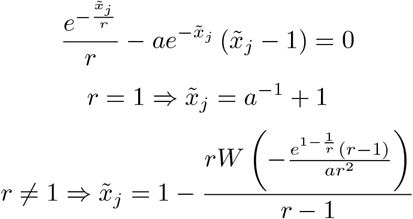

### 6.5 Simulations

We simulated *l* individuals 100 times with random initial locations inside a *d* dimensional torus of size *s* with scaled attraction *a* and scaled repulsion *r*. We used a torus to prevent edge effects of the simulation area and chose the size of the torus to be an order of magnitude larger than the effective interaction range of the individuals to limit the effects of the torus on the movement process. We simulated our model using Euler integration with a custom-build adaptive step size routine until all movements converged. We performed these simulations for all combinations of:

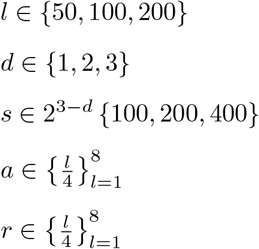

Given that in a *d*-dimensional torus there are 2^*d*^ straight paths between two points, we computed at each time step in the simulations the sum of all 2^*d*^ (*l* − 1) scaled net attraction vectors 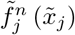 over toroidal distances 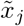 per individual *j*.

### 6.6 Clustering

Unfortunately it proved to be too error-sensitive to directly determine the number of clusters and their sizes for each converged simulation. Therefore we computed for each simulation the minimum toroidal Euclidian distance matrix *M* and averaged it out element-wise for the 100 iterations of each parameter combination of the simulation. Finally, we transformed *M* to 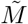 so that it quantifies the deviation from the expected distance to the *k*th neighbour under complete spatial randomness, by:

1. Sorting the rows per column in ascending order.

2. orting the columns per row in ascending order, to retrieve a matrix of columns with ascending distances to the *k*th neighbour.

3. Removing all columns larger than

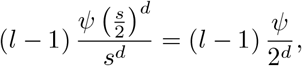

where *ψ* is the volume of a *d* dimensional ball with unit size

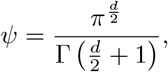

to include only the *k* neighbours that with complete spatial randomness are inside a *d* dimensional ball of diameter *s* within a *d* dimensional cube of size *s*.

4. Scaling every column by the expected average distance to the *k*th neighbour under complete spatial randomness [32]

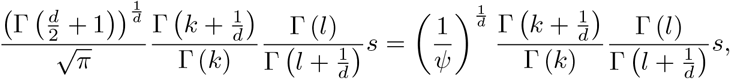

which (given that *l* is rather large) can be reduced with Stirling’s approximation to

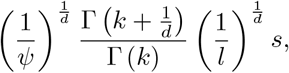

which can be approximated for large values of *k* with

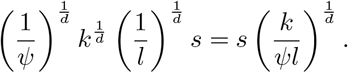

### 6.7 Point Process Model

From 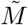 the number of clusters with a certain size were derived by identifying the matrix cells with a substantially lower value than 1 (see Figure 3, bottom row). After having identified these cells with a lower value, the probability distribution of each individual being in a cluster of a certain size could easily be determined. To verify that this procedure matched the actual group size distribution, we computed 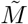 for a custom-build Point Process Model with known group sizes and (based on the known group sizes) marked the matrix cells that signified distances within a cluster (Figure 7). We performed this procedure with Point Process Models using all combinations of the following parameters:

1. Mean *µ* (*Ñ*) of the Beta distribution of the number of clusters *N*, where
2. Standard deviation *σ* (*Ñ*) of the Beta distribution of *N* relative to the maximum possible standard deviation (*µ* (1 − *µ*)), ∈ {0.5, 0.7, 0.9}
3. Concentration 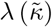 of the symmetric Dirichlet distribution of the number of individuals *κ* per cluster (with the number of categories of the distribution being fixed at *N* ), where 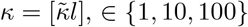, ∈ {1, 10, 100}
4. Repulsion coefficient *ω* of the clusters as a function of cluster size *κ*, where the location of all cluster centres *g* are simulated in a single *ABM* using 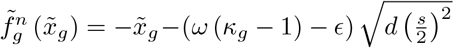, with 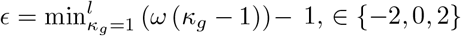
5. Inter-individual distance *θ* within each cluster, simulated with a separate *ABM* for each cluster using 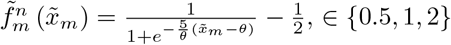
6. Number of individuals *l*, ∈ {50, 100, 200}
7. Number of dimensions *d*, ∈ {1, 2, 3}
8. Torus size *s*, ∈ 2^3−*d*^ {100, 200, 400}

**Figure 7.**
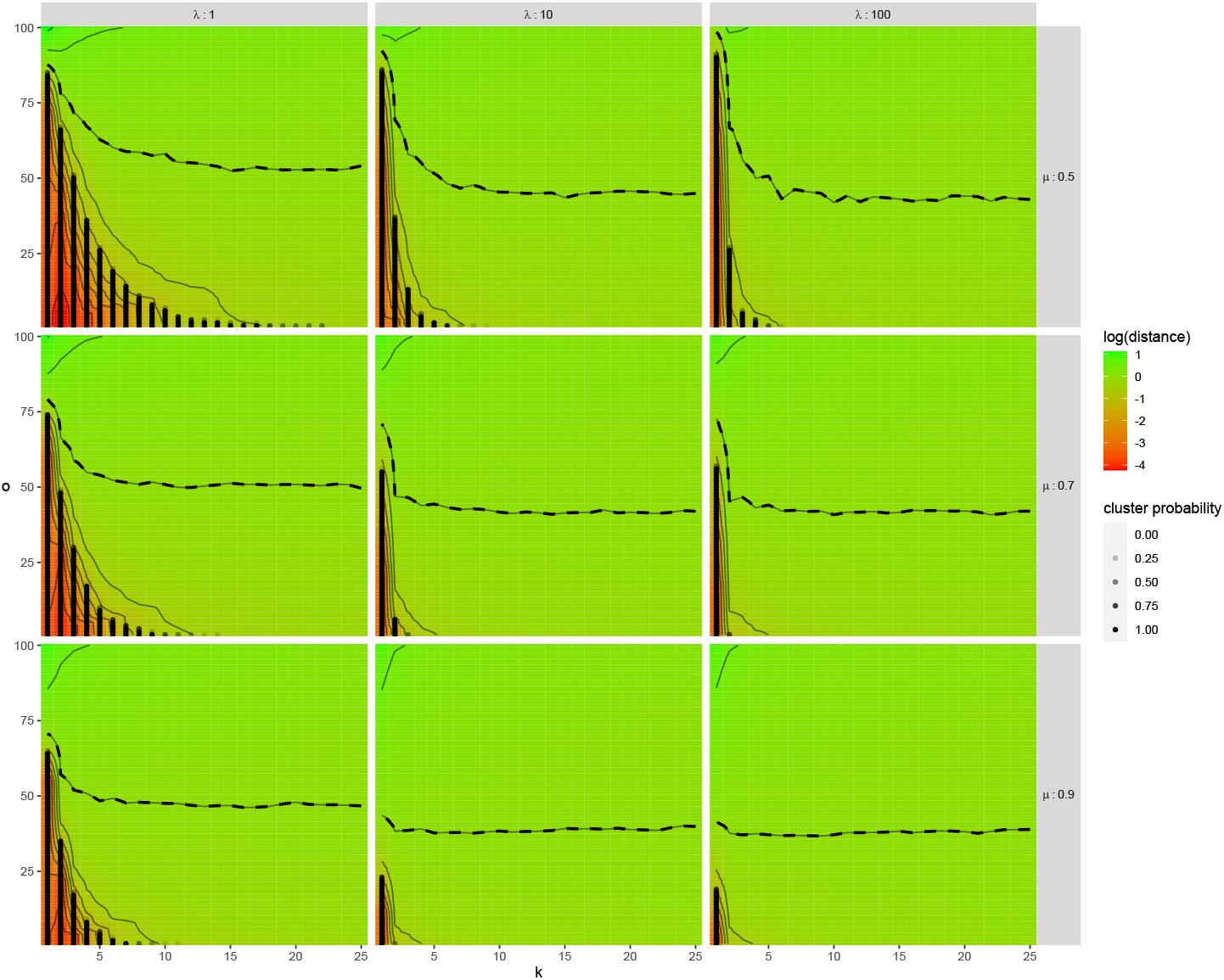
Average transformed distance matrices 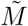 of the converged states of all 100 Point Process Model initializations, with the *k*th neighbour (until the 25th) on the x-axis, the ascending order *o* of distances on the y-axis, all values of group size concentration *λ* in the top panels and all values of mean number of clusters *µ* in the side panels. All other parameters of the model are set at the median value for this figure. The black dots signify the distances that are known to be from within a cluster, which matches accurately with the low values of 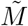.

## Data accessibility

The scripts and simulations are available to the reviewers upon request and will be uploaded to a public repository upon acceptance of the manuscript.

## Authors’ contributions

HdK, JE, AD and FvL developed the research aim; JE and HdK designed the methods; JE wrote the code and analysed the simulations; JE led the writing of the manuscript; all authors (JE, AD, FvL and HdK) contributed critically to the analyses and manuscript. All authors gave final approval for publication.

## Acknowledgments

We thank Tom Nijman and Muhammed Ercan for their work on preliminary analyses.

## References

[1] Nathan R, Getz WM, Revilla E, Holyoak M, Kadmon R, Saltz D, et al. A movement ecology paradigm for unifying organismal movement research. Proceedings of the National Academy of Sciences. 2008;105(49):19052– 19059.

[2] Brown JS, Laundré JW, Gurung M. The Ecology of Fear: Optimal Foraging, Game Theory, and Trophic Interactions. Journal of Mammalogy. 1999;80(2):385–399.

[3] Laundré JW, Hernández L, Altendorf KB. Wolves, elk, and bison: reestablishing the “landscape of fear” in Yellowstone National Park, U.S.A. Canadian Journal of Zoology. 2001;79(8):1401–1409.

[4] Lima SL. Back to the basics of anti-predatory vigilance: the group-size effect. Animal Behaviour. 1995;49(1):11–20.

[5] Krause J, Ruxton GD. Living in groups. New York, U.S.A.: Oxford University Press; 2002.

[6] Couzin ID, Krause J. Self-Organization and Collective Behavior in Vertebrates. In: Advances in the Study of Behavior. vol. 32. Academic Press; 2003. p. 1–75.

[7] Alexander RD. The Evolution of Social Behavior. Annual Review of Ecology and Systematics. 1974 11;5(1):325–383.

[8] Hamilton WD. Geometry for the selfish herd. Journal of Theoretical Biology. 1971;31(2):295–311.

[9] Isbell LA. Contest and scramble competition: patterns of female aggression and ranging behavior among primates. Behavioral Ecology. 1991;2(2):143– 155.

[10] Krause J, Godin JJ. Shoal Choice in the Banded Killifish (Fundulus diaphanus, Teleostei, Cyprinodontidae): Effects of Predation Risk, Fish Size, Species Composition and Size of Shoals. Ethology. 2010 4;98(2):128–136.

[11] Hoare DJ, Couzin ID, Godin JGJ, Krause J. Context-dependent group size choice in fish. Animal Behaviour. 2004 1;67(1):155–164.

[12] Couzin ID, Krause J, James R, Ruxton GD, Franks NR. Collective memory and spatial sorting in animal groups. Journal of Theoretical Biology. 2002;218(1):1–11.

[13] Sinclair ARE. The African Buffalo: A Study of Resource Limitation of Populations. Chicago, USA: University of Chicago Press; 1977.

[14] Fryxell JM, Mosser A, Sinclair ARE, Packer C. Group formation stabilizes predator–prey dynamics. Nature. 2007 10;449(7165):1041–1043.

[15] Sorato E, Gullett PR, Griffith SC, Russell AF. Effects of predation risk on foraging behaviour and group size: adaptations in a social cooperative species. Animal Behaviour. 2012 10;84(4):823–834.

[16] Brouwer L, Cockburn A, van de Pol M. Integrating Fitness Components Reveals That Survival Costs Outweigh Other Benefits and Costs of Group Living in Two Closely Related Species. The American Naturalist. 2020 2;195(2):201–215.

[17] Sciortino F, Mossa S, Zaccarelli E, Tartaglia P. Equilibrium Cluster Phases and Low-Density Arrested Disordered States: The Role of Short-Range Attraction and Long-Range Repulsion. Physical Review Letters. 2004 7;93(5):055701.

[18] Liu Y, Xi Y. Colloidal systems with a short-range attraction and long-range repulsion: Phase diagrams, structures, and dynamics. Current Opinion in Colloid & Interface Science. 2019 2;39:123–136.

[19] Gazi V, Passino KM. A class of attractions/repulsion functions for stable swarm aggregations. International Journal of Control. 2004;77(18):1567– 1579.

[20] Bernoff AJ, Topaz CM. A Primer of Swarm Equilibria. SIAM Journal on Applied Dynamical Systems. 2011;10(1):212–250.

[21] Leverentz AJ, Topaz CM, Bernoff AJ. Asymptotic Dynamics of Attractive-Repulsive Swarms. SIAM Journal on Applied Dynamical Systems. 2009;8(3):880–908.

[22] Eftimie R, de Vries G, Lewis MA, Lutscher F. Modeling Group Formation and Activity Patterns in Self-Organizing Collectives of Individuals. Bulletin of Mathematical Biology. 2007;69:1537–1565.

[23] Romanczuk P, Schimansky-Geier L. Swarming and pattern formation due to selective attraction and repulsion. Interface Focus. 2012 12;2(6):746–756.

[24] Chen Y, Kolokolnikov T. A minimal model of predator-swarm interactions. Journal of The Royal Society Interface. 2014;11:20131208.

[25] Niwa HS. School Size Statistics of Fish. Journal of Theoretical Biology. 1998 12;195(3):351–361.

[26] Thaker M, Vanak AT, Owen CR, Ogden MB, Slotow R. Group Dynamics of Zebra and Wildebeest in a Woodland Savanna: Effects of Predation Risk and Habitat Density. PLoS ONE. 2010 9;5(9):e12758.

[27] Ebensperger LA, Wallem PK. Grouping increases the ability of the social rodent, Octodon degus, to detect predators when using exposed microhabitats. Oikos. 2002 9;98(3):491–497.

[28] Gillespie T, Chapman C. Determinants of group size in the red colobus monkey ( Procolobus badius ): an evaluation of the generality of the ecological-constraints model. Behavioral Ecology and Sociobiology. 2001 9;50(4):329–338.

[29] Herbert-Read JE, Rosén E, Szorkovszky A, Ioannou CC, Rogell B, Perna A, et al. How predation shapes the social interaction rules of shoaling fish. Proceedings of the Royal Society B: Biological Sciences. 2017 8;284(1861):20171126.

[30] Frommen JG, Hiermes M, Bakker TCM. Disentangling the effects of group size and density on shoaling decisions of three-spined sticklebacks (Gasterosteus aculeatus). Behavioral Ecology and Sociobiology. 2009 6;63(8):1141–1148.

[31] Scott-Samuel NE, Holmes G, Baddeley R, Cuthill IC. Moving in groups: how density and unpredictable motion affect predation risk. Behavioral Ecology and Sociobiology. 2015 6;69(6):867–872.

[32] Bhattacharyya P, Chakrabarti BK. The mean distance to the n th neighbour in a uniform distribution of random points: an application of probability theory. European Journal of Physics. 2008;29(3):639–645.

